# Genome-wide association mapping of sudden death syndrome resistance in germplasm from the USDA soybean collection

**DOI:** 10.64898/2026.09.29.755259

**Authors:** Lilian. M. Miranda, Adam Mahan, Zhanyou Xu

## Abstract

Sudden Death Syndrome (SDS), caused by *Fusarium virguliforme*, can significantly impact soybean yields. To advance the understanding of SDS resistance in diverse plant introductions, we conducted genome-wide association studies (GWAS) on a set of 351 accessions from the USDA National Plant Germplasm System (NPGS) Soybean Germplasm Collection. The analysis revealed seven single nucleotide polymorphisms (SNPs) significantly associated with foliar SDS resistance; three significant QTL (on chromosomes 3, 9 and 11) were detected with Fixed and Random Model Circulating Probability Unification (FarmCPU) and four (on chromosomes 5, 6 and two on chromosome 7) with Bayesian-information and Linkage disequilibrium Iterative Nested Keyway (BLINK). Localized linkage disequilibrium blocks were estimated for these regions. Notably, all of them overlapped with regions where QTL associated with SDS resistance had been previously reported and were within, or close to, sequences annotated as implicated in plant defense mechanisms, such as toxin transporters and transcription factors. The architecture of SDS resistance appears quantitative and polygenic, with most loci explaining only modest phenotypic variance. Six plant introductions in this study—PI 175188, PI 307889A, PI 323559, PI 578307C, PI 445683, and PI 323558— harbor favorable alleles at all seven loci, providing high-value resources for resistance introgression. This study underscores the value of leveraging the USDA Soybean Germplasm Collection for the discovery of resistance loci and provides a foundation for marker-assisted breeding to enhance SDS resistance in elite soybean lines.

## Introduction

Sudden Death Syndrome (SDS), caused by the soil-borne fungus *Fusarium virguliforme* O’Donnell & Aoki, is a major threat to soybean production, resulting in significant yield losses across diverse environments [1]. The pathogen infects tap roots and lower stems, causing vascular discoloration and, during mid to late reproductive stages, producing foliar symptoms such as interveinal chlorosis and necrosis that lead to early defoliation and pod abortion. These above-ground symptoms are attributed to toxins produced by the fungus and transported via the xylem [2].

Advances in quantitative trait locus (QTL) mapping have revealed several loci associated with SDS resistance. A review by [3] introduced the nomenclature *Rfv* (resistance to *Fusarium virguliforme*) for ten consistently reported loci, each documented in at least three independent studies. Notably among them, are the QTL on chromosomes 17 and 18 [4, 5]. The SDS resistance QTL on chromosome 17 was identified in both PI 567374 and the cultivar Ripley through analyses conducted in two independent mapping populations [5]. The chromosome 18 QTL has demonstrated the largest effect in some populations, explaining up to 36.4% of resistance variation [4], although its effectiveness can be highly context-dependent and, in certain genetic backgrounds, the putatively resistant allele has been associated with increased disease severity [6]. Epistatic interactions between QTLs, such as those on chromosomes 8 and 18, have been shown to significantly increase the proportion of phenotypic variance explained, justifying a shift toward more sophisticated breeding models [7].

The complexity of SDS resistance is further underscored by the weak correlation between root and foliar symptoms, each governed by distinct genetic loci [8]. Moreover, the synergistic relationship between SDS and soybean cyst nematode (SCN) further complicates disease management. This relationship was demonstrated in a study of at-planting risk factors, which found that SCN egg quantity was the primary factor explaining variation in end-of-season SDS severity, and that final SDS levels, in turn, accounted for most of the variation in soybean yield [9]. A meta-analysis estimated a yield decrease of 0.5% for every unit increase in the foliar disease index [1].

Integrated management strategies, including seed treatments, increased planting densities, extended crop rotations, and host resistance—are essential for mitigating yield losses [10–12]. However, reliance on fungicides with single-site modes of action (fluopyram and pydiflumetofen) raises concerns about the rapid development of fungicide resistance [12]. Genomic selection for SDS resistance has shown limited prediction accuracy, likely due to low heritability and poor genetic correlations between traits [13].

SDS resistance is rare, with most of the USDA Soybean Germplasm accessions evaluated for this disease exhibiting susceptibility [14, 15]. Recent large-scale screening efforts have identified over 100 plant introductions with foliar disease severity equal to or lower than the resistant check PI 567374, offering options for new sources of resistance [16]. However, the practical use of these sources is often constrained by the negative yield linkage drag associated with exotic germplasm. The identification of molecular markers associated with resistance loci will enable targeted introgression of specific genomic regions and is therefore critical for a more efficient introgression of SDS resistance into elite soybean lines.

This research aims to advance our understanding of the genetic architecture of SDS resistance by conducting genome-wide association mapping in a diverse population of soybean accessions that includes these recently phenotyped plant introductions, thereby making these resources more useful and informative for breeders and researchers.

## Materials and Methods

### Plant Materials

The genome wide association analysis included the 125 plant introductions evaluated in at least 4 replicates (most of them rated as resistant to foliar disease severity) by [16] and 226 additional entries from the NPGS Germplasm Resources Information Network (GRIN) (https://npgsweb.ars-grin.gov/gringlobal/search) selected because they have a numerical score for SDS and have been genotyped with the Illumina Infinium SoySNP50K BeadChip [17].

### Phenotypic data

*Fusarium virguliforme* isolate Mont-1 was used for all the evaluations. The inoculation procedure was described by [18] and [16]. All values were converted to % severity using the mid-point value from the scale by Chawla et al. (2013). In order to combine the two data sets, the 125 plant introductions evaluated by [16] were scaled to the GRIN data set using the common checks as reference points. An adjustment factor was determined by calculating the absolute arithmetic difference between the mean of the checks in GRIN and the mean of the checks for the 125 plant introductions. The common checks used were PI 567374, PI 536636 (cv. Ripley), and PI 525454 (cv. Spencer). Finally, we performed rank-based inverse normal transformation [19] using SAS version 9.3 (SAS institute, Cary, NC).

### Genotypic data

The genotypic data for all entries was obtained from the SoySNP50K BeadChip [17] available in soybase (www.soybase.org). Missing SNPs were imputed using Beagle-geno version 5.4 [20] and SNPs with minor allele frequency <5% were removed from the analysis. The final number of SNP markers included in the analysis was 33,691.

### Association Analyses

PLINK 2.0 was used to calculate principal component (PC) eigen values and eigen vectors that were later used to graph principal components vs percent of variance explained in R. Genome wide association analysis was conducted using the GAPIT R package (Lipka et al. 2012). Three different models; Mixed Linear Model (MLM), Bayesian-information and Linkage disequilibrium Iterative Nested Keyway (BLINK) and Fixed and Random Model Circulating Probability Unification (FarmCPU); were used for the analysis. A standalone ordinary least squares (OLS) linear regression was calculated to confirm a significant QTL with very low minor allele frequency using R. The linkage disequilibrium (LD) decay was estimated for a 500 kb window on each significant chromosome region using PLINK 1.9 [21]. The LD matrices (pairwise, squared allele frequency correlations between SNPs) generated by PLINK 1.9 were plotted in R to generate LD heat maps. The LD blocks (r^2^ > 0.6) containing the SNP peaks were searched for candidate genes using the soybean reference genome Wm82.a4.v1 available in Soybase (www.soybase.org).

## Results

The PC analysis revealed significant population stratification. The curve of PC eigen values versus % variation explained (Fig 1) flattened after four PCs, therefore four PCs were used in the GAPIT analysis. No significant QTL were detected with MLM. With an LOD threshold of 6, three significant regions were detected with Farm CPU and four with BLINK (Table 1, Fig 2). There was no significant overlap between the significant hits detected by the two methodologies, with the possible exception of the significant QTL on chromosome 5 detected with BLINK that was very close to the significant threshold in the FarmCPU results. Despite this, the Q-Q plots for both Farm CPU and BLINK demonstrated robust control over population structure, evidenced by the observed values tightly tracking the null distribution line. Sharp, late-stage upward deflections for both models indicate high-confidence, true genetic associations (Fig 3). The cumulative phenotypic variance explained (PVE) by the multi-locus model frameworks ranged from 4.4% (peak SNP ss715590926 on chromosome 5) to 39.8% (peak SNP ss715598157 on chromosome 7). Because of the unusually high PVE value estimated for chromosome 7, an estimate of the marginal phenotypic variance explained (R²) by this locus was calculated using a standalone ordinary least squares (OLS) linear regression. The single-marker model confirmed a highly significant association between marker ss715598157 and SDS severity (p = 6.53 e-05) but the estimated value for PVE was only 4.47%.

**Fig 1.**
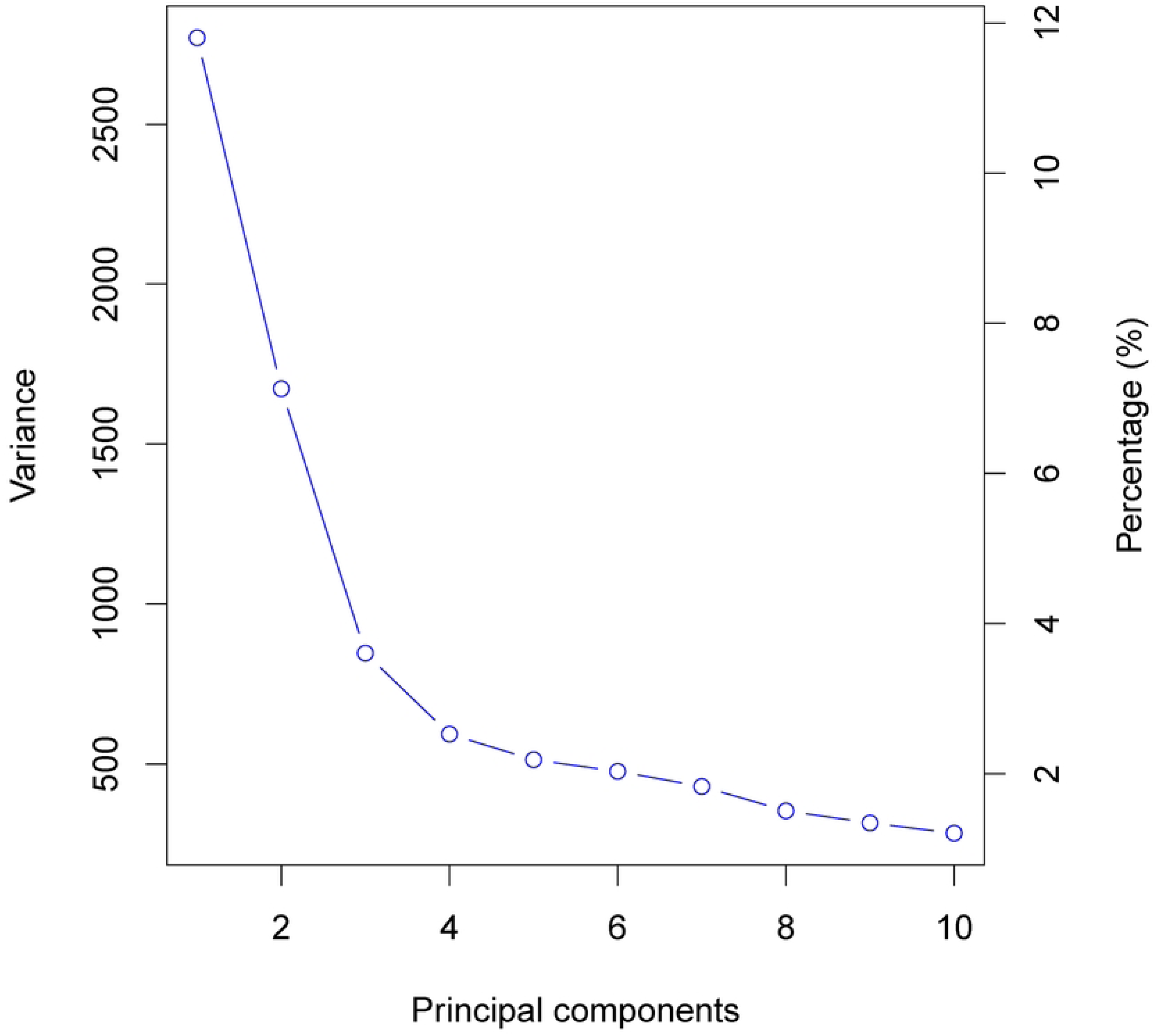
Principal Components versus percentage of variance explained for sudden death syndrome severity in a population of 351 soybean accessions.

**Fig 2.**
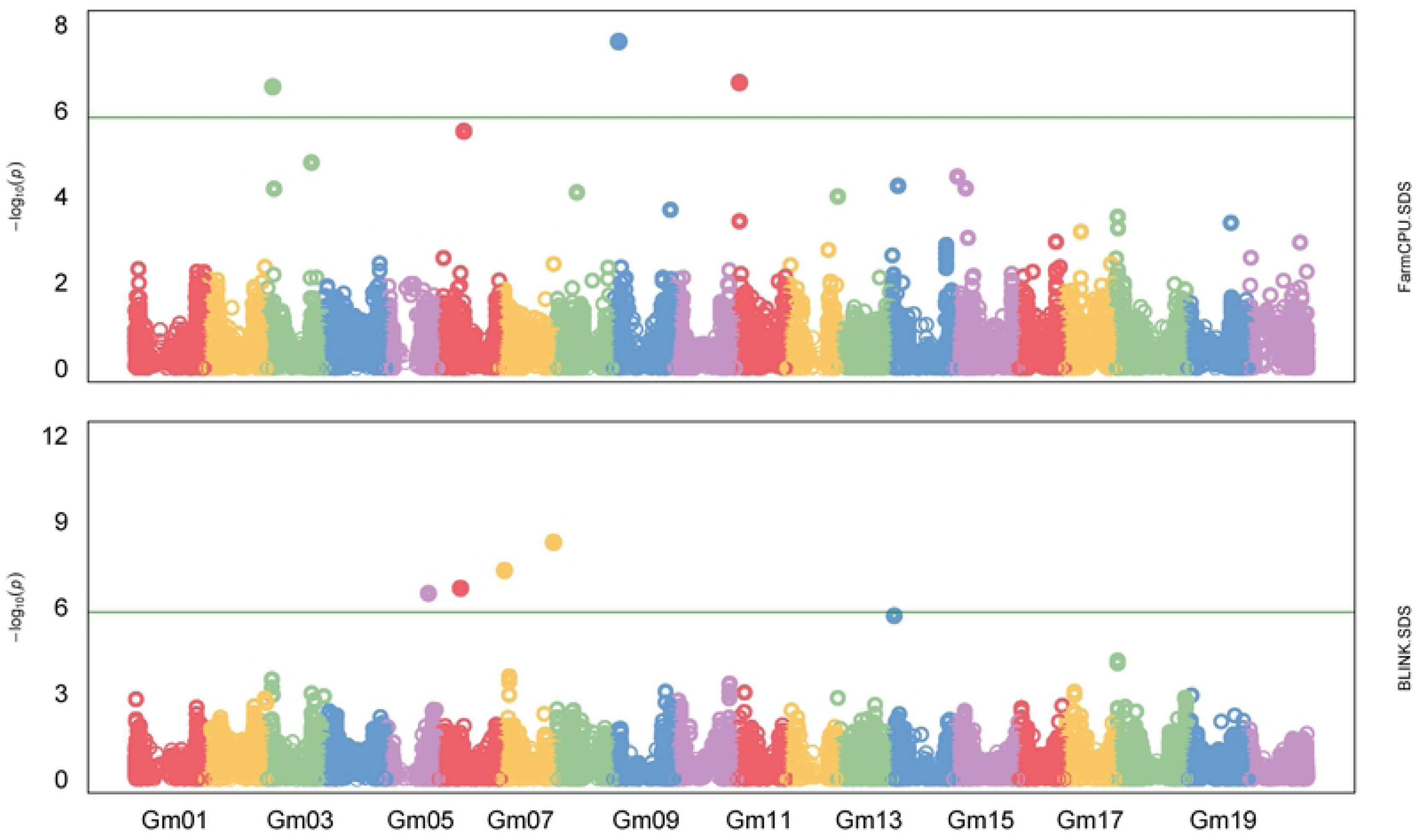
Manhattan plots of GWAS for sudden death dyndrome foliar disease severity. The upper plot corresponds to the Fixed and Random Model Circulating Probability Unification model (FarmCPU), and the lower plot is for the Bayesian-information and Linkage disequilibrium Iterative Nested Keyway (BLINK) model. The x-axis of each Manhattan plot represents the chromosome number, and the y-axis represents the LOD (-log10(p-value)).

**Fig 3.**
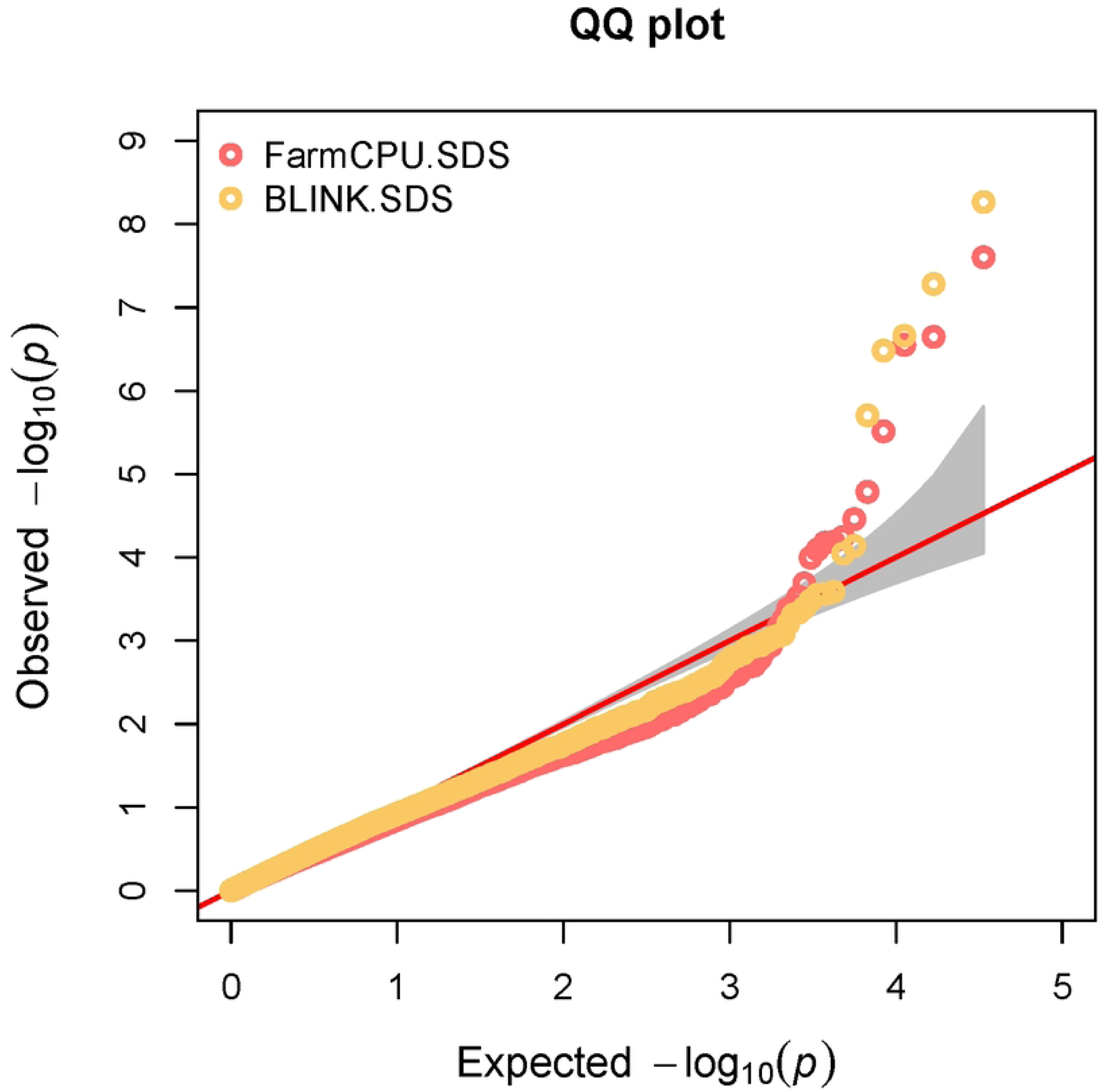
Quantile-quantile plot for the Fixed and Random Model Circulating Probability Unification (Farm CPU) and the Bayesian-information and the Linkage disequilibrium Iterative Nested Keyway (BLINK) models. The x-axis of the QQ-plot represents the expected -log10(p-value) and the y-axis displays the observed -log10(p-value).

**Table 1.**
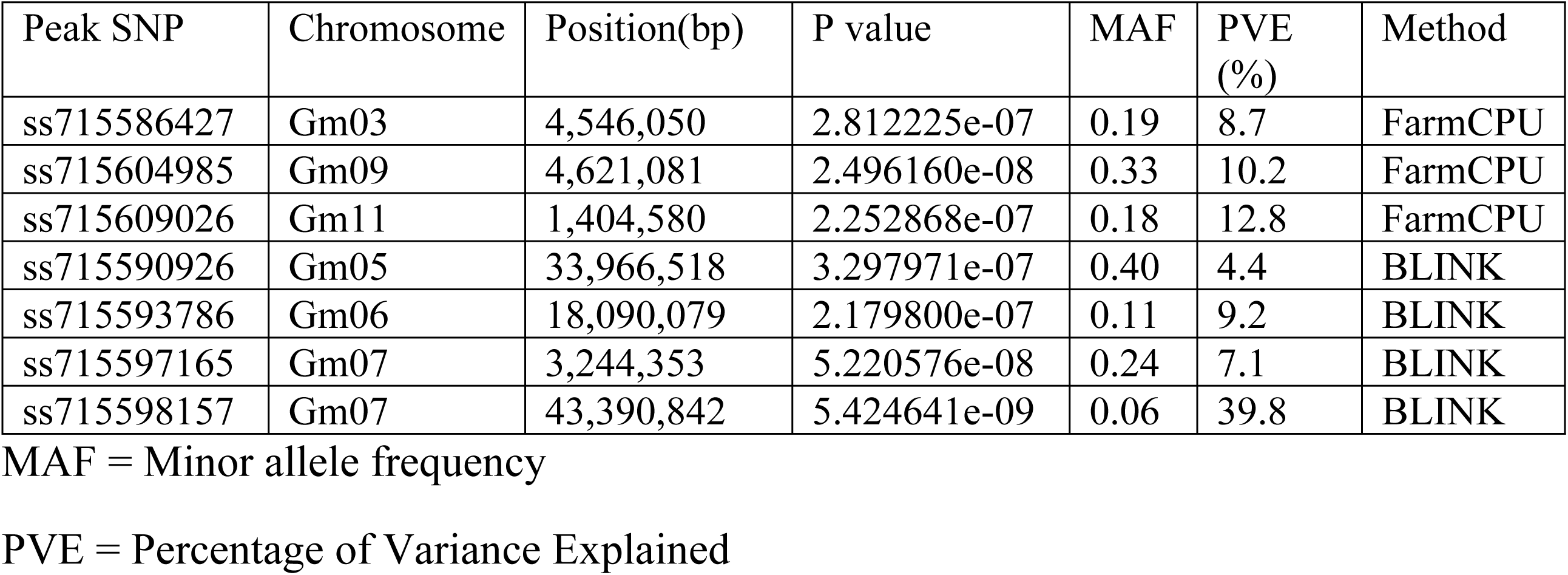
Significant QTL detected in the Genome Wide Association Analysis of 351 plant introductions from the USDA Soybean Germplasm Collection.

The localized linkage disequilibrium decay estimated around each significant hit showed variable linkage patterns on each chromosome (Figures 4-10) which were subsequently used to determine the boundaries of the candidate gene search windows. Candidate genes identified within each chromosomal window are shown in Table 2.

**Fig 4.**
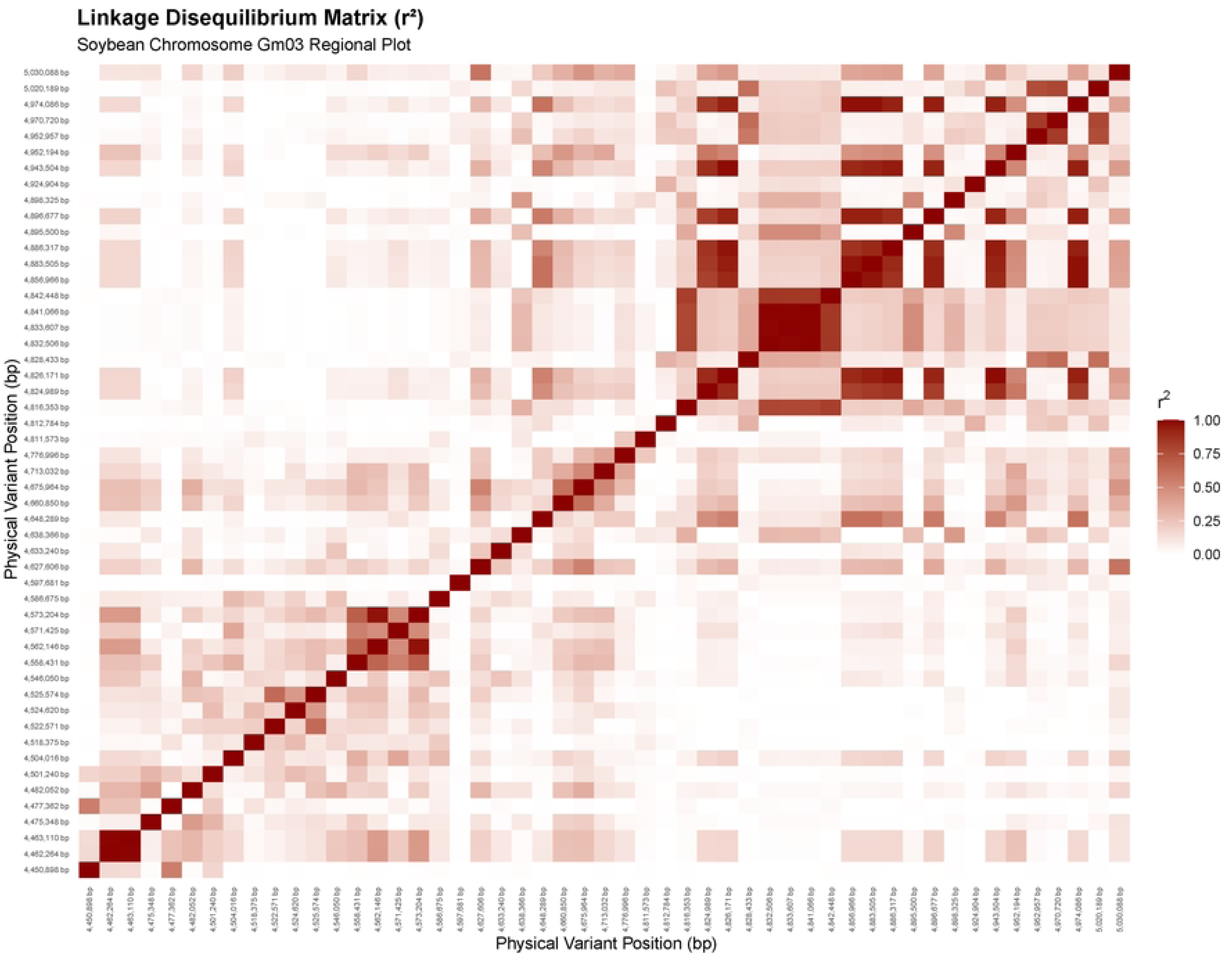
Heat map of the linkage disequilibrium matrix for the region surrounding the significant SNP peak ss715581174 on Chromosome 3. X and y axes display SNP positions in bp.

**Fig 5.**
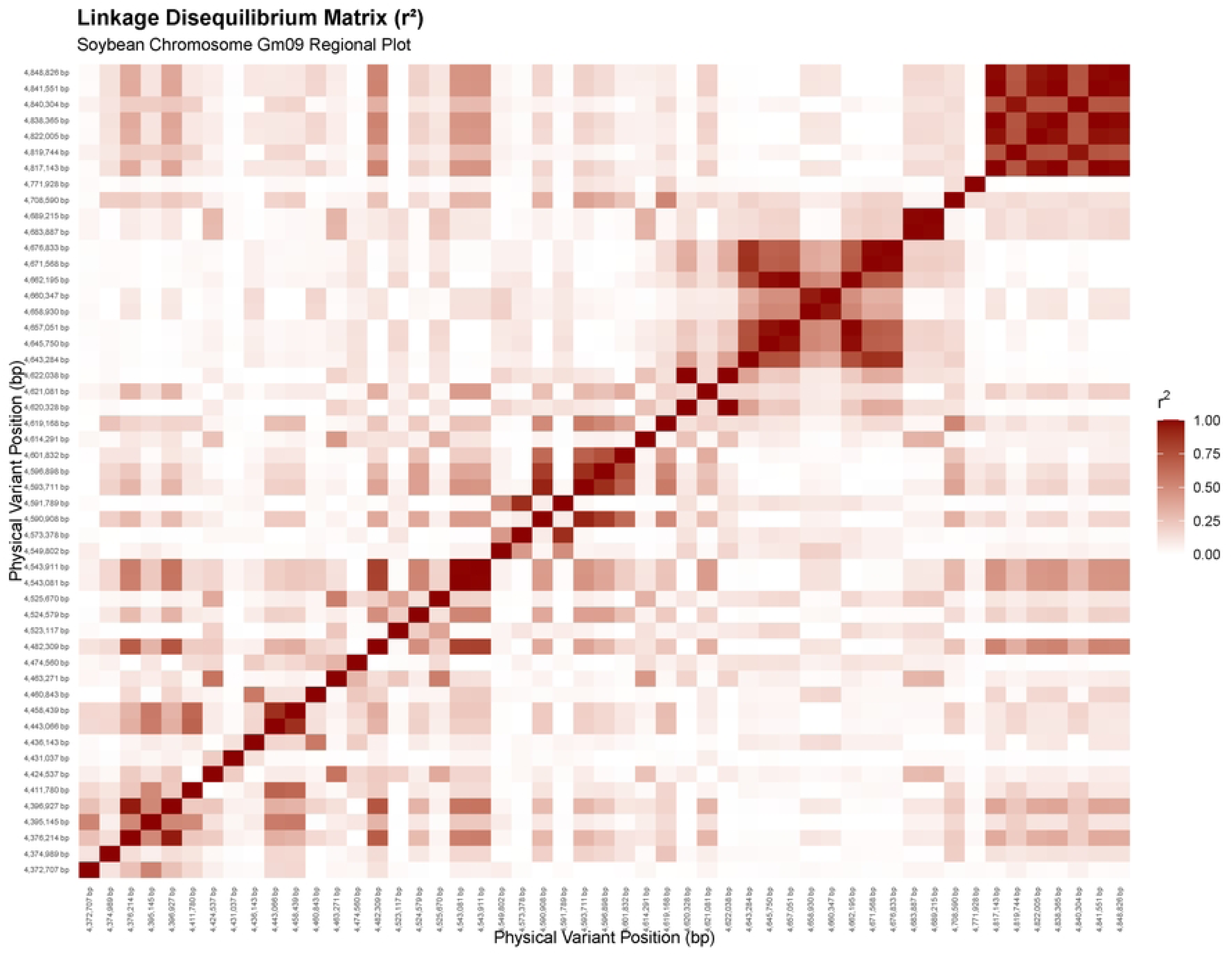
Heat map of the linkage disequilibrium matrix for the region surrounding the significant SNP peak ss715560495 on Chromosome 9. X and y axes display SNP positions in bp.

**Fig 6.**
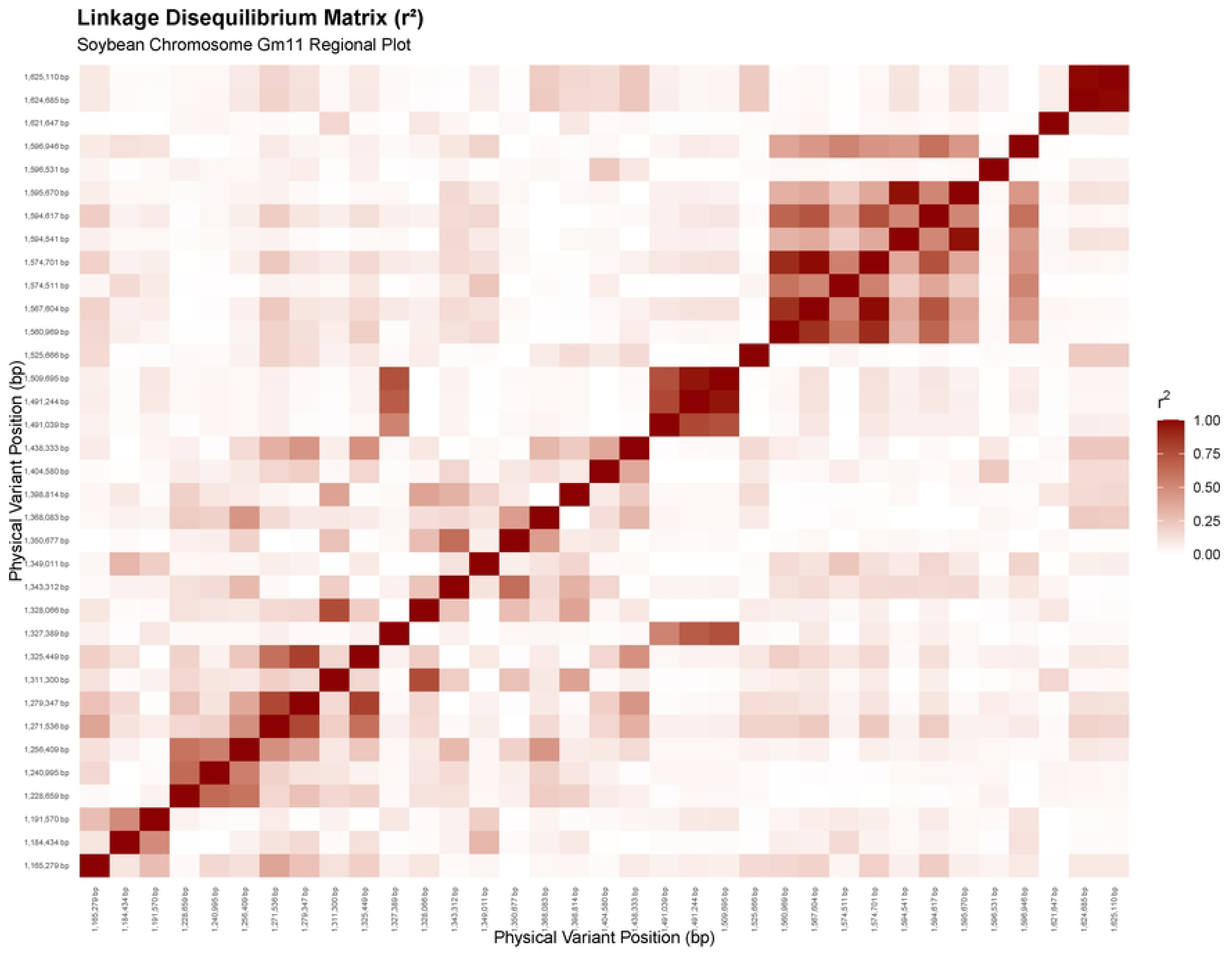
Heat map of the linkage disequilibrium matrix for the region surrounding the significant SNP peak ss715609026 on Chromosome 1. X and y axes display SNP positions in bp.

**Table 2.** Functional prediction of candidate genes in the interval of significant chromosomal positions associated with resistance to Sudden Death Syndrome.

| Candidate Gene | Chromosome | SNP Peak Position | Coordinates | Functional Description |
| --- | --- | --- | --- | --- |
| Glyma.03G036200 | 3 | 4,546,050 bp | 4,546,504 to 4,553,151bp | MATE efflux family protein |
| Glyma.05G145000 | 5 | 33,966,518 bp | 33,932,216 to 33,938,685 bp | ATP-binding ABC transporter |
| Glyma.06G198100 | 6 | 18,090,079 bp | 17,950,985 to 17,954,921 bp | Flavin-binding monooxygenase family protein |
| Glyma.07G039800 | 7 | 3,244,353 bp | 3,278,455 to 3,281,813 bp | Cytochrome P450 superfamily protein |
| Glyma.07G252832 | 7 | 4,339,0842 bp | 43,337,069 to 43,339,066 bp | MFS transporter |
| Glyma.09G053000<br>Glyma.09G052800<br>Glyma.09G052900 | 9 | 4,621,081 bp | 4,635,186 to 4,618,301bp | Ethylene-responsive transcription factors |
| Glyma.09G053700<br>Glyma.09G054000<br>Glyma.09G054600<br>Glyma.09G054200<br>Glyma.09G054100 | 9 | 4,621,081 bp | 4,685,986 to 4,814,070 bp | Ankyrin repeat-containing protein |
| Glyma.11G020200 | 11 | 1,404,580 bp | 1,413,431 to 1,418,274 bp | Calcium-dependent protein kinase 6 |

On chromosome 3, the peak SNP captured by FarmCPU (ss715581174) resides within a ∼124kb block spanning from 4.46 Mb to 4.59 Mb (Table 2 and Fig 4). This peak marker is only 454 bp away from a gene encoding a MATE (multi drug and toxic compound extrusion) transporter. On chromosome 9 the local Linkage Disequilibrium (LD) architecture surrounding the significant FarmCPU marker peak (ss715560495 at 4,621,081 bp) placed the hit between 4.62Mb and 4.71Mb (Table 2 and Fig 5). A cluster of three Ethylene Responsive Transcription Factors (ERFs) is located 2 to 10 kb from the SNP peak and a gene encoding an ankyrin repeat containing protein resides within the same block. The significant FarmCPU hit on Chromosome 11 (ss715609026 at 1,404,580 bp) was positioned within an LD block from 1.33Mb to 1.44Mb (Table 2 and Fig 6) with a Calcium-Dependent Protein Kinase 6 (CDPK6) located just 8.8kb away from the SNP peak.

For the BLINK models, the significant SNP peak ss715590926 on chromosome 5 (33,966,518bp) was positioned in an LD window between 33.91Mb and 33.98Mb (Table 2, Fig 7), which contains an ATP-Binding cassette (ABC) transporter gene. On chromosome 6, the significant BLINK variant ss715593786 (18,090,079 bp) anchors a linkage block spanning from 17.7Mb to 18.1Mb (Table 2, Fig 8) that encompasses a gene encoding a Flavin-Binding Monooxygenase (FMO) family protein. The BLINK model also identified a peak variant at position 3,244,353 bp on chromosome 7 (region 1), marking the upstream boundary of a distinct LD block from 3.2 Mb to 3.3 Mb (Table 2, Fig. 9), which contains a cytochrome P450 superfamily protein gene. Finally, an independent significant region on chromosome 7 (region 2) was anchored by the BLINK marker ss715598157 (43,390,082 bp), defining the downstream boundary of a separate high-LD block spanning 43.3 Mb to 43.4 Mb (Table 2, Fig. 10), within which a major facilitator superfamily (MFS) transporter gene is located.

**Fig 7.**
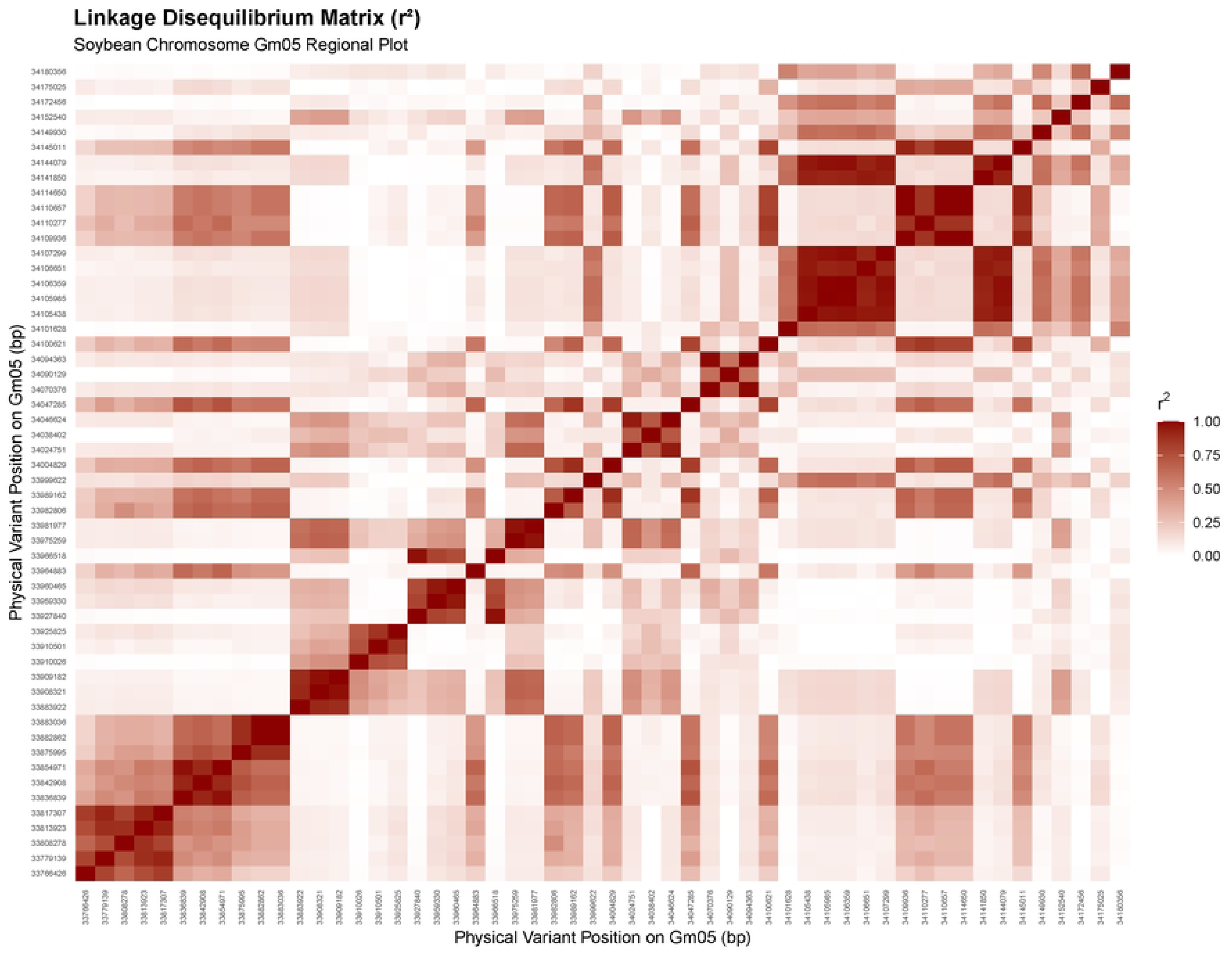
Linkage disequilibrium matrix plot for the region surrounding the Significant SNP peak ss715590926 on Chromosome 5. X and y axes display SNP positions in bp.

**Fig 8.**
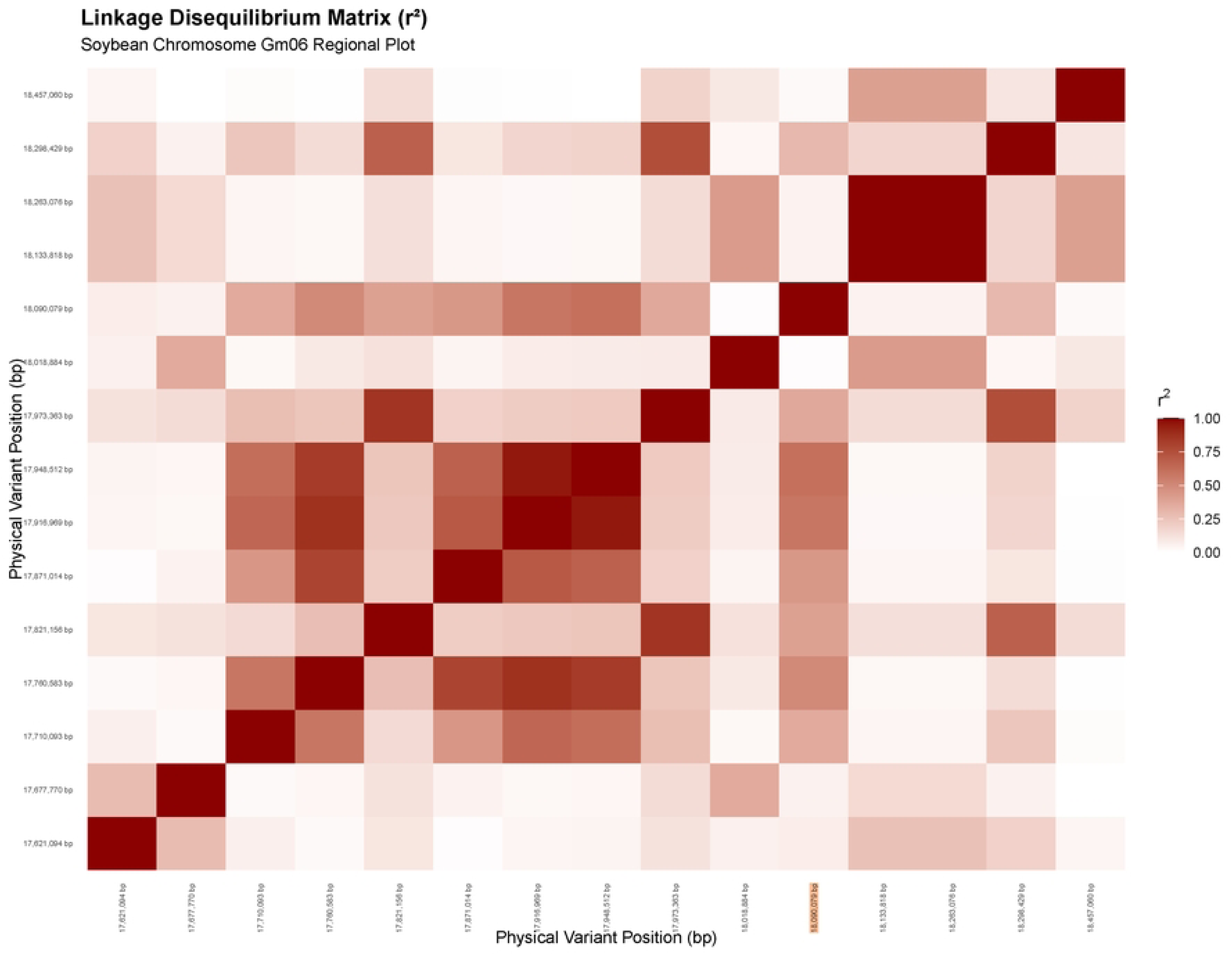
Heat map of the linkage disequilibrium matrix for the region surrounding the significant SNP peak ss715593786 on Chromosome 6. X and y axes display SNP positions in bp.

**Fig 9.**
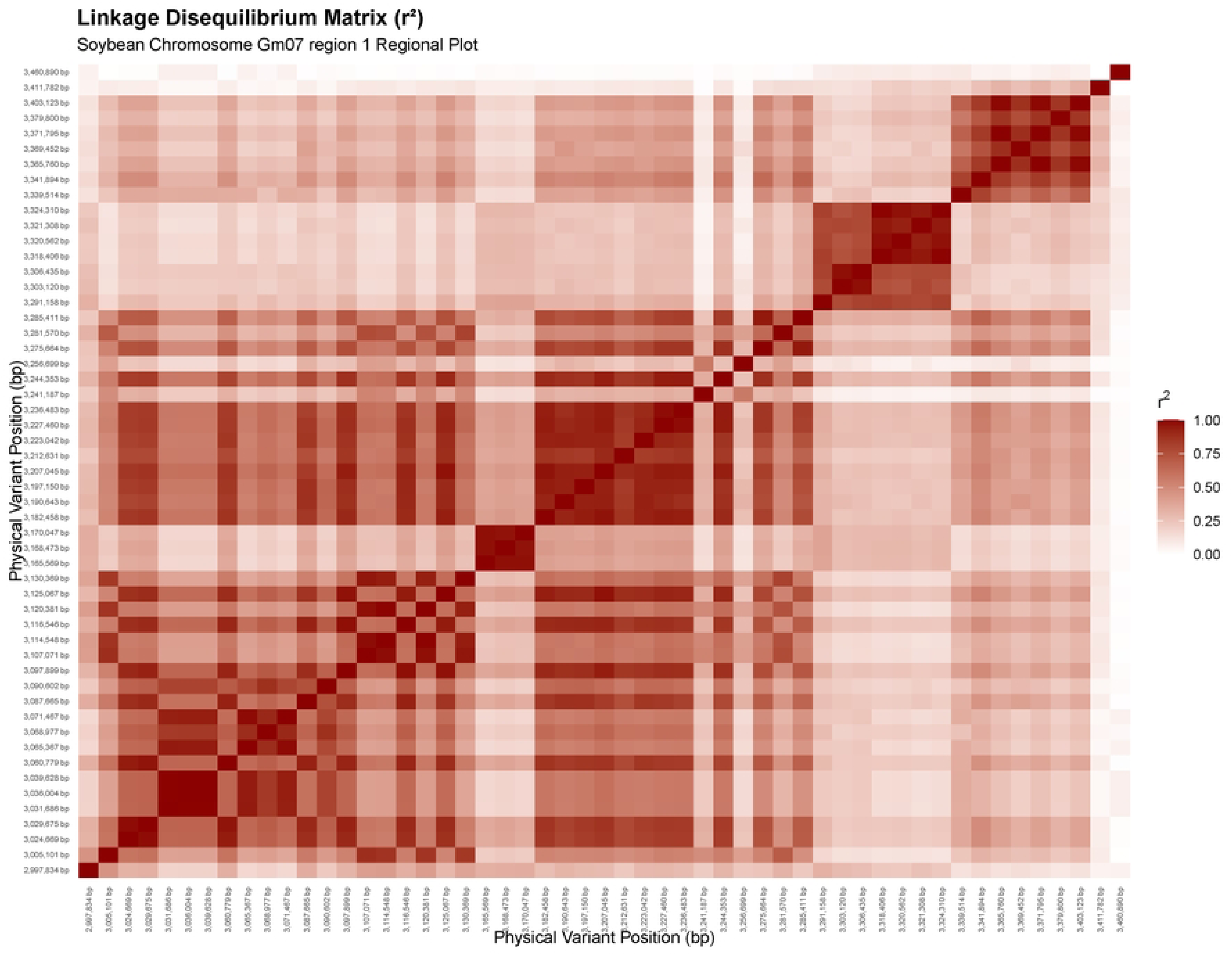
Heat map of the linkage disequilibrium matrix for the region surrounding the significant SNP peak ss715597165 on Chromosome 7. X and y axes display SNP positions in bp.

**Fig 10.**
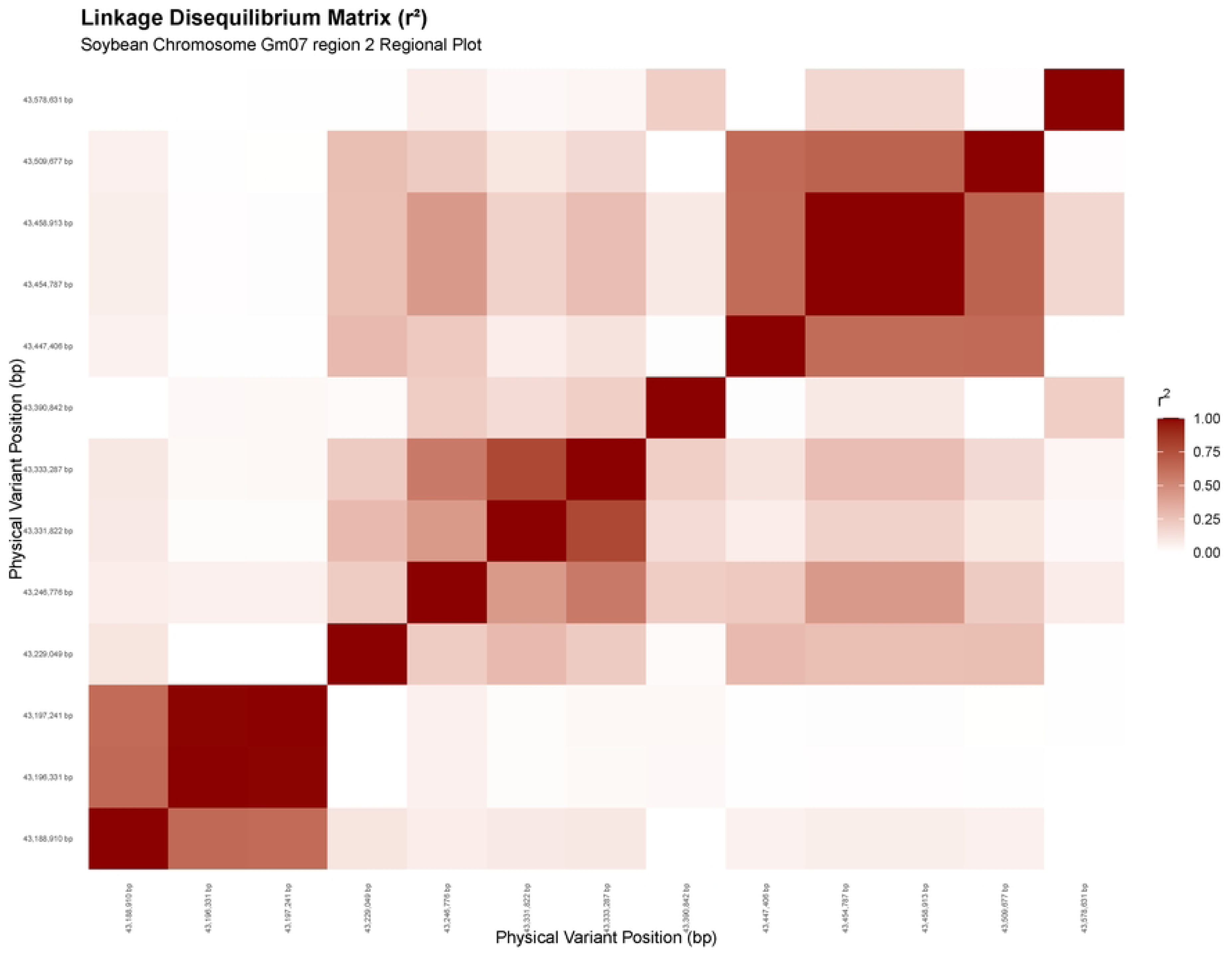
Heat map of the linkage disequilibrium matrix for the region surrounding the significant SNP peak ss715598157 on Chromosome 7. X and y axes display SNP positions in bp.

## Discussion

The GWAS analysis detected 7 significant QTL associated with SDS resistance. The lack of overlap between the BLINK and FarmCPU results may be explained, in part, by the differences in the algorithms deployed in each methodology. BLINK uses a localized, fixed-effect approach guided by the Bayesian Information Criterion (BIC), allowing it to focus on small linkage disequilibrium (LD) blocks and efficiently isolate narrow peaks. FarmCPU, in contrast, applies a fixed genome-wide optimization window within a global kinship matrix. Differences in the way these models account for background genetic variation and marker effects can therefore result in the identification of different loci. Consequently, associations corresponding to relatively narrow or low-frequency haplotype blocks may be detected by BLINK but not by FarmCPU.

Although the genetic marker ss715598157 on chromosome 7 explained only 4.47% of the total variation in SDS resistance when analyzed alone using ordinary least squares (OLS) regression, the cumulative multi-locus BLINK model accounted for 39.8% of the phenotypic variance. Several factors may contribute to this discrepancy. First, the favorable allele was relatively rare in the population studied (minor allele frequency = 0.06), which limits the proportion of population level phenotypic variance that can be attributed to this marker when evaluated individually. Moreover, the single marker OLS estimate may be affected by population structure and therefore should not be interpreted as an absolute estimate of the biological contribution of the locus. A more precise estimation of the individual effect of this locus would require a mapping population with more balanced allele frequencies and controlled genetic backgrounds such as a bi-parental population. Despite these population specific limitations, the likelihood of this locus representing a true SDS-resistance QTL is strongly supported by previous reports of SDS resistance loci in the same region [7, 22]. The relatively small proportion of phenotypic variance explained (PVE) by individual loci is also consistent with previous studies demonstrating the complex and quantitative genetic architecture of SDS resistance in soybean [3, 7, 8].

Due to the autogamous nature of soybeans, whole-genome linkage disequilibrium (LD) decay typically extends up to about 300 kb. However, the patterns of LD blocks can differ considerably between euchromatic and heterochromatic regions. Rather than applying a uniform genome-wide physical window, candidate genes in the present study were identified using a locus-specific approach based on the boundaries of local haplotype blocks surrounding each significant SNP. This approach provided a more restrictive and biologically informed search interval while accounting for local variation in LD. Even within these sometimes-narrower intervals, candidate genes with known or putative roles in plant defense and stress responses were identified for all significant SNPs.

On chromosome 3, the close proximity of the peak SNP ss715581174 to a MATE transporter gene suggests a likely role in disease resistance; MATE transporters serve as vital components of the plant defense response by mediating the active efflux of fungal toxins [23]. Similarly, a cluster of three ethylene responsive transcription factors (ERFs) was situated 2 to 10 kb from the peak variant detected by FarmCPU on chromosome 9. Ethylene is an important regulator of defense responses against soil borne necrotrophic and hemi biotrophic fungal pathogens [24]. This same LD region harbors genes encoding ankyrin repeat containing proteins that regulate plant signaling networks during stress responses [25]. The chromosome 11 significant locus resides only 8.8 kb from a Calcium-Dependent Protein Kinase 6 (CDPK6). CDPKs have been associated with downstream defense-related signaling cascades in soybeans [26] and other crops [13].

The BLINK model also identified additional loci within LD blocks containing genes encoding defense-related transport and metabolic proteins. The candidate window on chromosome 5 contains an ATP-binding cassette transporter; ABC proteins actively export fungal phytotoxins, xenobiotics, and antimicrobial compounds out of the host plant cell to neutralize cellular damage during pathogen attacks [27]. On chromosome 6, the candidate window captures a Flavin-binding monooxygenase (FMO) family gene, which encodes a class of key enzymes responsible for triggering Systemic Acquired Resistance (SAR) signals in legumes [28]. The upstream locus identified by BLINK on chromosome 7 resides within an LD region that contains a cytochrome P450 super family gene. Cytochrome P450 enzymes have been previously associated with resistance to *Phytophtora sojae* [29]. Lastly, the significant chromosome 7 marker ss715598157 anchors a linkage block containing a Major Facilitator Superfamily (MFS) transporter gene. MFS are widely hypothesized to be involved in the targeted export of fungal pathogens toxins out of host cells [30].

The proximity of these defense-associated genes to the significant SNPs supports their consideration as candidate genes, although functional validation will be required to determine whether they contribute directly to SDS resistance. Taken together, the functional diversity of these candidate genes; including transport, signaling, and metabolism; provides further support for the multigenic nature of SDS resistance and illustrates the potential of multi-locus GWAS approaches to resolve components of the genetic architecture underlying complex disease resistance.

The SNP peaks detected on Chromosome 3 (ss715586427) and Chromosome 6 (ss715593786) align with the intervals for the loci designated as *Rfv03-1* (43.96-45.78Mb)and *Rfv06-02* (16.33-23.85 Mb) respectively by [3]. These loci were designated based on their consistent identification in at least three independent studies, supporting their reproducibility as genomic regions associated with SDS resistance. The remaining significant SNPs on Chromosomes 5, 7, 9, and 11 also fall within genomic regions containing previously reported quantitative trait locus (QTL) for sudden death syndrome (SDS) resistance in soybean, as documented by [7, 22, 31].

The major QTL on chromosome 17 (*Rfs1*) [5] and chromosome 18 (*rhg1/Rfs2*) [32] were not detected in this population. Although resistant alleles at these loci are present in some of the germplasm included in this study, including Ripley and PI 567374, their low frequencies may have limited the power of this GWAS analysis to detect their effects. This observation underscores an important limitation of association mapping in diverse germplasm like the USDA soybean germplasm collection: rare resistant alleles are difficult to detect because the small number of individuals carrying them provided limited power. Consequently, GWAS should be complemented by developing bi-parental populations, which allow for mapping alleles with more balanced frequencies and can also be implemented and used in parallel within breeding programs.

Six plant introductions in this study—PI 175188, PI 307889A, PI 323559, PI 578307C, PI 445683, and PI 323558—carry all seven favorable alleles associated with resistance to SDS at the significant SNPs identified by GWAS. According to [16] four of these lines (PI 175188, PI 307889A, PI 578307C, and PI 445683) exhibited foliar disease severity scores of less than 1.5 on a scale of 1 to 8 [18] during step VII of a tiered testing sequence. This final tier included 12 replications and followed earlier evaluations with one replication in tiers I–V and four replications in tier VI. The remaining two plant introductions, PI 323559 and PI 323558, were both classified as resistant to foliar symptoms in the study by [33]. In the same study, PI 323559 also showed resistance to root rot. The presence of the seven favorable alleles in these accessions, together with their consistently reported resistance phenotypes, identifies these plant introductions as promising sources of SDS-resistance alleles for marker-assisted breeding and introgression.

## Conclusions

The QTL identified in this study expands the genomic resources available for SDS resistance and provides additional markers for evaluating and deploying resistance alleles from the USDA Soybean Germplasm Collection. In particular, the identification of multiple favorable alleles within highly resistant plant introductions provides opportunities to combine resistance loci through marker-assisted selection. Further validation of these loci across genetic backgrounds and environments, together with functional characterization of the candidate genes, will help determine their individual contributions to SDS resistance and facilitate their incorporation into soybean breeding programs.

## Notes

### Competing Interest Statement

The authors have declared no competing interest.

